# Characterization of genetically novel chimeric plasmids from *Salmonella* Heidelberg conferring multi-drug resistance and increased pathogenicity

**DOI:** 10.1101/2025.10.27.684768

**Authors:** Jonathan Betts, Andreas Liaropoulos, Matteo Buffoni, Michael S. M. Brouwer, Kees T. Veldman, Francisco J. Salguero, Roberto La Ragione, Daniela Ceccarelli, Apostolos Liakopoulos

## Abstract

*Salmonella enterica* serotype Heidelberg (*S*. Heidelberg) is a significant cause of human salmonellosis, with resistance to extended-spectrum cephalosporins posing challenges for clinical management. In this study, we genetically and functionally characterized six plasmids isolated from distinct PFGE-types of *S*. Heidelberg previously reported in The Netherlands, revealing a chimeric IncC-I1a plasmid that shares similarities with the epidemic pESI plasmid identified in emergent *S*. Infantis isolates. Our analysis showed the accumulation of genetic determinants conferring resistance to antibiotics (*bla*_CMY-2_, *tetA* and *sul2*) and heavy metals (*mer* operon), as well as enhanced pathogenicity (Yersinia high-pathogenicity island). All these elements are located on a stable non-conjugative but likely mobilizable plasmid backbone that imposes only a marginal fitness cost on its bacterial host. This convergence of multidrug resistance and pathogenicity likely enhances bacterial adaptability and virulence, undermining control strategies based solely on reducing selective pressure. The emergence and dissemination of such hybrid plasmids represent an increasing threat to both public and animal health, analogous to that posed by pESI-like plasmids, and underscore the urgent need for integrated genomic surveillance and risk assessment, as their continued expansion could complicate antimicrobial therapy and containment efforts during *Salmonella* outbreaks.

## Introduction

*Salmonella enterica* serotype Heidelberg (*S*. Heidelberg) is one of the most predominant serotypes causing human salmonellosis in Canada and the United States, and it has increasingly been reported in Europe^1, 2, 3^. It is frequently isolated from poultry and poultry meat and has been associated with multiple food related outbreaks^4^. Notably, it is resistant to extended-spectrum cephalosporins (ESCs), which are essential for treating systemic *S*. Heidelberg infections in children and in adults when fluoroquinolones are contraindicated^5, 6^. In addition to resistance to these first-line antibiotics, *S*. Heidelberg carries a diverse array of virulence-associated determinants^4^, which contribute to its pathogenicity and increase the risk of invasive infections^5, 7^. The potential for this serotype to acquire further virulence factors heightens public health concerns and underscores the need for monitoring of its evolving pathogenic potential.

Although not yet identified in S. Heidelberg, the chimeric virulence plasmid pESI (IncI1 and IncP-1α) and its genetic variants have drawn significant attention for their role in the global dissemination of *Salmonella enterica* serovar Infantis. These chimeric pESI-like megaplasmids encode not only several virulence factors, such as the iron acquisition system yersiniabactin and the chaperone-usher fimbriae Ipf and Klf, but also genes conferring antibiotic resistance and stress tolerance^9^, posing major challenges for both clinical management and public health. Given the established pathogenicity and antibiotic resistance profile of *S*. Heidelberg, the potential acquisition of such plasmids could further enhance its fitness and complicate prevention and control efforts.

In the present study, we report the genetic and functional characterization of chimeric IncC-I1a megaplasmids recovered from *S*. Heidelberg strains genetically reminiscent of pESI-like plasmids, documenting the emergence of plasmids encoding antibiotic and heavy metal resistance, and increased pathogenicity within this *Salmonella* serotype. Given that the acquisition of these plasmids may further promote the spread of *S*. Heidelberg, potentially following a trajectory similar to that of *S*. Infantis., our findings underscore the importance of genomic surveillance in detecting emerging plasmid-mediated threats among this high-risk *Salmonella* serotype.

## Materials and methods

### Bacterial isolates, transformants and plasmids

Six previously reported IncA/C plasmids encoding *bla*_CMY-2_ recovered during the national antimicrobial resistance monitoring program in the Netherlands were included in this study for further analysis. These plasmids were isolated from *S*. Heidelberg isolates belonging to distinct PFGE-types (XbaI.1965, XbaI.1973 and XbaI.X)^3^. *E. coli* DH10b transformants harbouring each of these plasmids, were generated as previously described^3^, and included in this study. Relevant characteristics of the specific plasmids are shown in Table S1. Due to the high synteny observed among these plasmids, a representative plasmid isolated from the predominant *Salmonella* PFGE-type (XbaI.1973) was randomly selected for further functional analysis.

### Plasmid sequencing, assembly and analysis

Plasmid DNA from transformants was isolated using the QIAfilter Plasmid Midi Kit (QIAGEN, Hilden, Germany) according to the manufacturer’s recommendations. Deep sequencing of the plasmid genomes was performed using 300-bp paired-end sequencing libraries prepared with the Nextera TAGmentation kit on an Illumina MiSeq platform. Long-read libraries were prepared with the Native barcoding kit EXP-NBD-103 and the Ligation Sequencing kit SQK-LSK108, and sequenced on a MinION Mk1B with an R9.4 flow cell. Subsequently, high-quality filtered Illumina- and Oxford Nanopore-derived reads were used to create *de novo* hybrid assemblies with Unicycler v0.4.7^10^. Putative open reading frames (ORFs) were identified by RAST version 2.0^11^ and subjected to BLASTP analyses against the NCBI non-redundant proteins (NR) database. ResFinder (version 3.0), PlasmidFinder (version 1.3), pMLST (version 1.4) and ISfinder were used to determine the presence of resistance genes, replicon types, plasmid sequence type and insertion sequences, respectively^12, 13, 14^. Further IncC subtyping was performed based on conserved backbone features, as previously described^15^. Relative pairwise distances of plasmid sequences were performed as previously described^16^, and the generated MUMi distance matrix was visualized using interactive tree of life (iTOL)^17^. Pairwise protein alignments between the three plasmid FASTA sequences were performed using the Sequence Alignment module of the PlasmidScope online tool^18^, which utilizes BLASTP to generate coverage and identity data. Custom R scripts, employing the *circlize*^19^ (for figure 2) or *genoPlotR*^20^ (for figure S1) packages, subsequently processed this alignment data, filtering for regions with at least 90% query coverage.

### Susceptibility testing assays

Minimum inhibitory concentrations (MICs) for the DH10b transformant harbouring the pSH-363.78 plasmid (DH10b::pSH-363.78) were determined in triplicate by broth microdilution, following the International Standard Organization 20776–1:2006 guideline. These assays aimed solely to confirm the resistance phenotype associated with the determinants identified on the novel IncC-I1a chimeric plasmids, therefore, the tested agents included were cefotaxime, sulfamethoxazole, tetracycline and mercuric chloride (HgCl_2_).

### Conjugal transfer assays

Plasmid conjugation was assessed in aerobic solid mating assays at 37 °C, using the chloramphenicol resistant (chlor^R^) *E. coli* MG1655::*yfp* as a recipient and DH10b::pSH-363.78 as donor in 1:1 ratio, as previously described^16^. The selection was made on LB agar supplemented with chloramphenicol (32 mg/L) and cefotaxime (1 mg/L).

### Fitness cost and stability assays

The growth rate for *E. coli* DH10b::pSH-363.78 was determined by a Synergy H1 plate reader (BioTek Inc., VT, USA) and measuring optical density at 600 nm every 15 min, as previously described^16^. Assays were performed in triplicate. Relative growth rate was calculated based on the generation time of the wild-type DH10b strain and student’s t-test was used to determine statistically significant differences, with a *P*-value ≤0.05 considered significant.

Stability of pSH-363.78 without selective pressure in an *E. coli* population was determined for 5 days (∼90 generations) in triplicate, as previously described^16^. Plasmid presence was confirmed by colony PCR using previously published primers targeting the *repA* gene of the IncA/C plasmids^21^.

### *Galleria mellonella* survival assays

*G. mellonella* larvae (TruLarv, Exeter, UK), in the final-instar larval stage were stored at 8°C before being randomly assigned to experimental groups. An inoculum test with 10^3^ to 10^6^ CFU/larvae was performed to determine the optimum bacterial inoculum for larval killing (LD_50_; approx. 50% mortality of larvae at 96 h post infection). Overnight cultures (16 h) of *E. coli* strains DH10b and DH10b::pSH-363.78 in LB broth were washed in phosphate buffered saline (PBS) before being serially diluted in PBS. Colony forming units (CFU) were determined by culturing the dilutions on LB agar and incubating aerobically at 37 °C for 24 h. Ten *G. mellonella* were each infected with 10 µl culture equating to 10^5^ CFU/larvae (LD50) into the hemocoels of the caterpillars via a left proleg injection using 25-μL Hamilton syringes (Cole-Parmer, London, United Kingdom). Larvae were incubated at 37 °C and scored for survival (live/dead) at 0, 24, 48, 72 and 96 h post infection, as previously described^22^. Phenotypic melanization scores for larvae were, also, recorded over 72 h as an indicator of morbidity based on a method previously described^23^. A score of 4 indicated total melanization of the larvae, 2 equated to melanin spots over the larvae, 1 equalled discoloration of the tail and a score of 0 represented no melanization, as previously described^24^.

### *Galleria mellonella* histopathology

*G. mellonella* larvae (3 per group) were injected with *E. coli* (DH10b or DH10b::pSH-363.78) at 10^6^ CFU/larvae or with a PBS control. Six hours post infection, larvae were fixed by immersion in 101% (v/v) neutral buffered formalin (NBF) for 14 to 21 days. Fixed larvae were bisected longitudinally to divide the body into two halves. At sectioning, an imprint of the haemolymph was taken into a glass slide, smeared, air dried and stained using the Gram-Twort method^25^. The tissue samples were embedded routinely in paraffin wax and 4-micron sections were stained with Haematoxylin and Eosin for microscopic examination, as previously described^25^. Semiquantitative scores for bacteria within the gut, bacteria within adipose bodies, internal melanin and haemocyte clusters were undertaken based on those previously described^25^.

### Accession number(s)

The nucleotide sequences of the reported plasmids have been deposited in GenBank under the following accession numbers CP080423.1 (pSH-369.05), CP080424.1 (pSH-366.65), CP080425.1 (pSH-363.78), CP080426.1 (pSH-365.07), CP080427.1 (pSH-359.42) and CP080428.1 (pSH-353.62).

## Results and discussion

### IncC plasmids of chimeric architecture

The six plasmids included in this study were initially classified as IncA/C using PCR-based replicon typing. However, whole-genome sequencing subsequently identified them as IncC plasmids belonging to sequence type (ST) 2 and possibly to backbone type 1b, although definitive classification as type 1b was not possible due to the complete absence of region 2 (R2), which is required for accurate subtyping^15^. These plasmids ranged in size from 209,580 bp (pSH-363.78) to 214,427 bp (pSH-369.05), with an average GC content of 51.8%. Nucleotide sequence analysis predicted between 231 ORFs in pSH-353.62 and 235 ORFs in both pSH-359.42 and pSH-366.65.

A comparison with 54 IncC plasmids from GenBank previously assigned to type 1b^15^ revealed that these six new plasmids possess a highly conserved and syntenic backbone (Figure S1), clustering together and distinctly from the rest of the known type 1b plasmids (Figure 1). The plasmid that clustered most closely with the plasmids characterized in this study was pCFSAN001921 (CP006050.1), previously identified in *S. enterica* serotype Typhimurium isolated from retail chicken breast meat^26^. Given the high sequence identity and synteny among these IncC plasmids, pSH-363.78 was selected as a representative for comparison with pCFSAN001921. The analysis revealed a deletion of approximately 57-kb in our plasmids relative to pCFSAN001921, encompassing the conserved IncC backbone region between the *traW* and *int* genes (Figure 2). Notably, this deletion coincided with the acquisition of an approximately 45-kb fragment flanked by IS21 and IS3 elements, which shares 99% sequence identity with segments of pSA02DT10168701_99, an IncI1α plasmid (CP012923.1) previously recovered from *Salmonella* Heidelberg isolated from chicken meat^27^. This plasmid belongs to ST12, which predominates among *S*. Heidelberg^3^. This suggests that the co-circulation of IncI1α-pST12 and IncC plasmids within this *Salmonella* serotype allowed for recombination events giving rise to the chimeric genetic architecture we observed.

**Figure 1.**
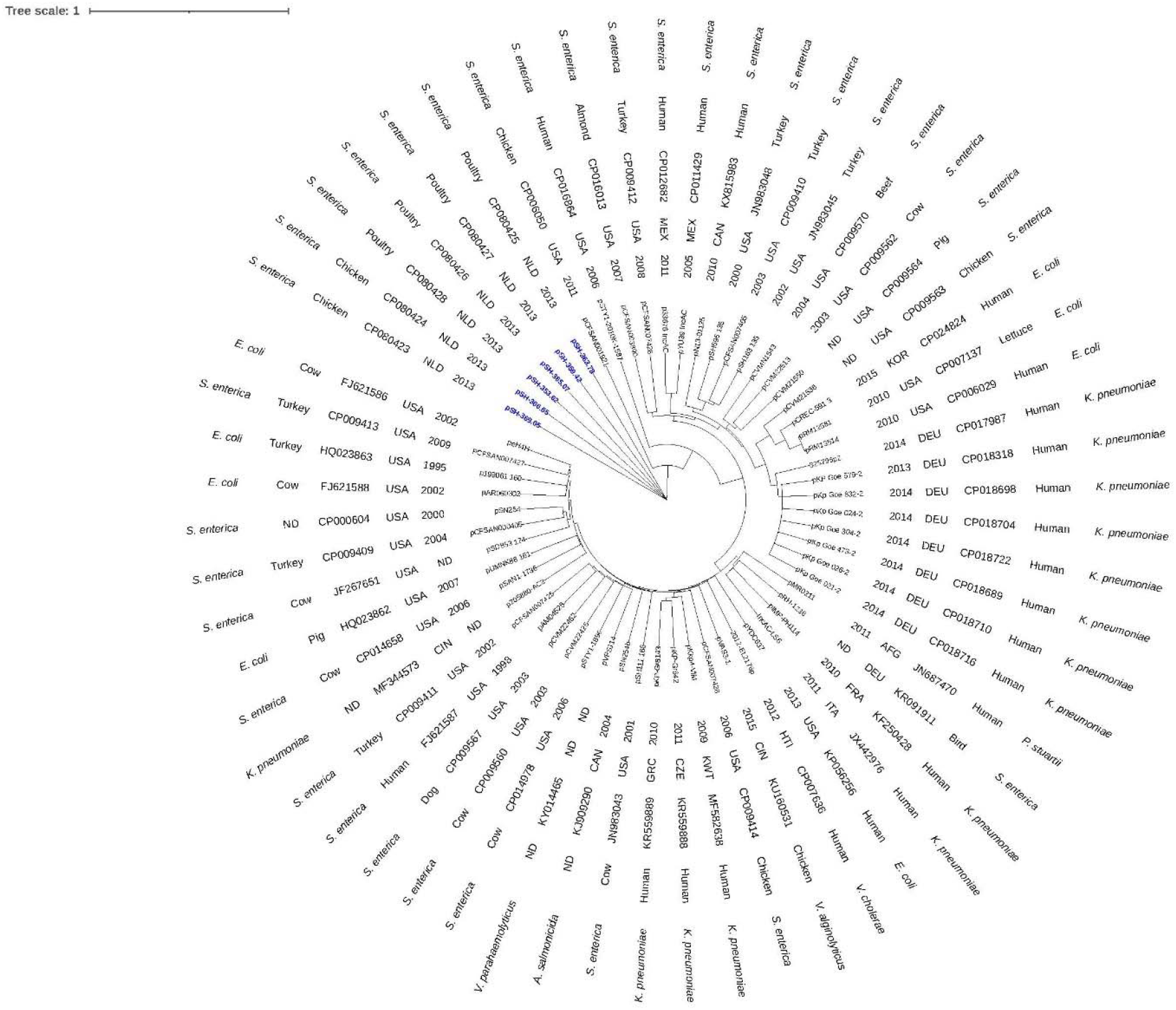
BioNJ phylogram based on MUMi distances among IncC type 1b plasmids. Plasmid sequences obtained in this study (indicated in blue) and those available in the GenBank database were compared pairwise, and maximum unique matches were converted to MUMi distances. The resulting distances were hierarchically clustered and visualized as a phylogram using the BioNJ algorithm using iTOL. Information on plasmid ID, year and country of isolation, GenBank accession number, sample origin, and bacterial host species are shown in this order. NLD: Netherlands; USA: United States of America; MEX: Mexico; CAN: Canada; KOR: South Korea; DEU: Germany; AFG: Afghanistan; FRA: France; ITA: Italy; HTI: Haiti; KWT: Kuwait; CZE: Czech Republic; GRC: Greece; CHN: China; ND: Not determined.

**Figure 2.**
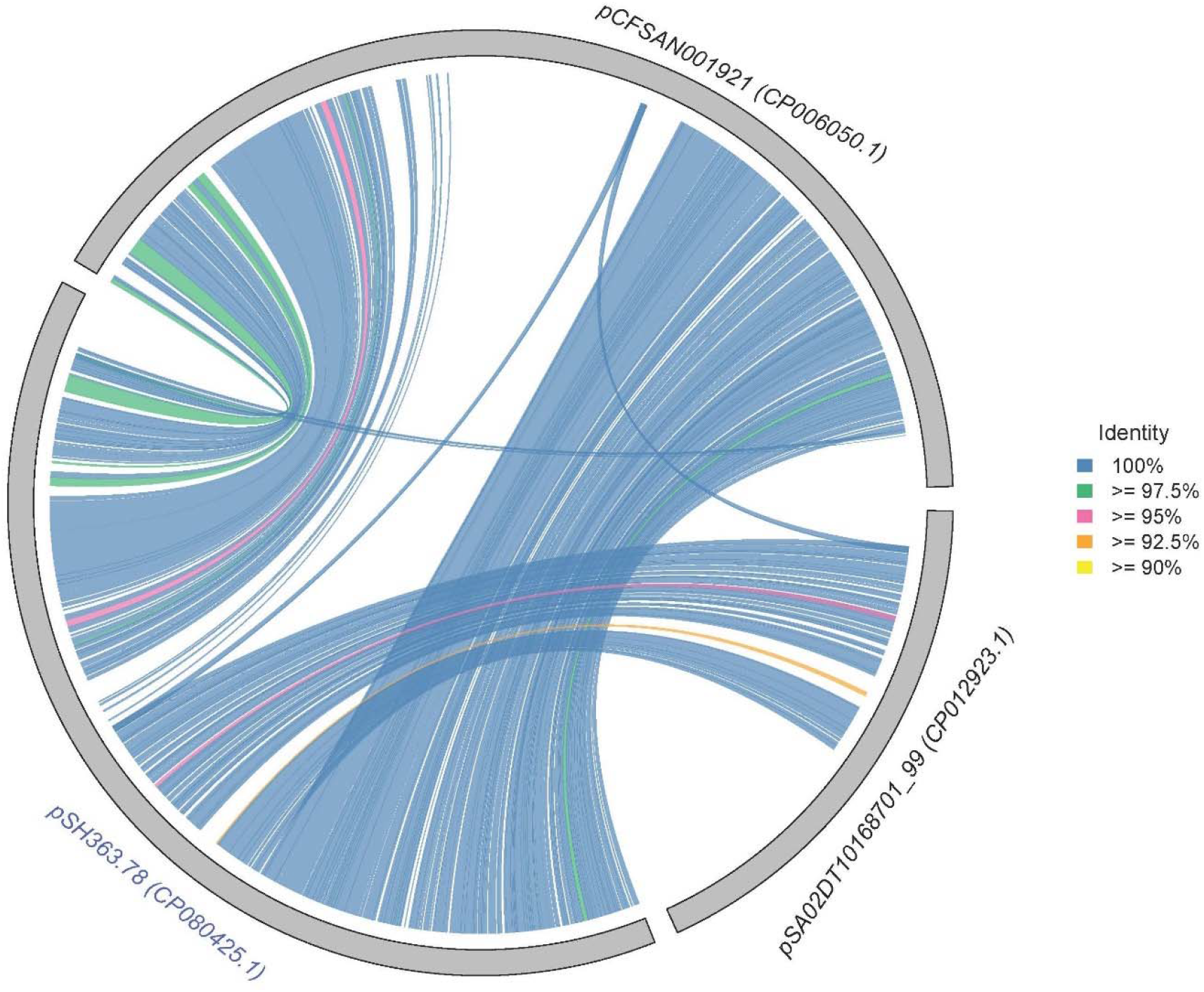
Circular synteny plot of homologous regions depicting sequence similarity among the novel chimeric IncC-I1a plasmid pSH363.78 (CP080425.1) and plasmids pCFSAN001921 (CP006050.1; IncC1b) and pSA02DT10168701_99 (CP012923.1; IncI1a-pST12), previously isolated from chicken meat. Each segment of the outer grey ring represents a complete plasmid genome and the inner coloured ribbons connect homologous coding regions between any two plasmids. Ribbon colour indicates the percentage identity of the corresponding protein alignments, as defined in the legend. Only pairwise protein alignments with at least 90% query coverage are displayed.

### Chimeric IncC-I1a plasmids encode resistance to multiple antibiotics and mercury

All six IncC-I1a plasmids were found to encode multiple antibiotic resistance determinants. Each plasmid carried the *bla*_CMY-2_, which confers resistance to extended-spectrum cephalosporins (*i*.*e*., cefotaxime MIC = 16 mg/L), embedded within the *ISEcp-1* tnpA-*bla*_CMY-2_-*blc*-*sugE* element spanning nucleotides 123,175 to 125,585 (nt numbering according to pSH-363.78). They also harboured the tetA efflux pump and its transcriptional repressor tetR between 42,044 and 43,999 nt, conferring resistance to tetracycline (MIC = 32 mg/L). This region was flanked by truncated *Tn3 tnpA* genes, located 1,449 bp upstream and 419 bp downstream. In addition, all plasmids carried the *sul2* gene between 45,420 and 46,235 nt, which confers resistance to sulfamethoxazole (MIC = 2048 mg/L). This gene was situated 564 bp upstream of the truncated *Tn3 tnpA* element that lies downstream of the *tetAR* operon. The co-localization of these resistance genes within a plasmid backbone enriched in mobile elements such as *ISEcp1* and truncated *Tn3* suggests ongoing recombination and genetic plasticity. This is consistent with previous reports describing the dynamic mosaic structure of IncC plasmids^15, 28^, thereby enabling the acquisition of additional resistance genes. Beyond these antibiotics and quaternary ammonium compounds resistance genes, all six plasmids also contained a multigene cluster (*mer* operon; 32,742 to 40,594 nt) conferring mercury resistance (MIC = 20 mg/L) that indicates potential co-selection under heavy metal exposure, as described in domestic animal-associated isolates^29^. Together, these findings highlight the evolutionary adaptability of these IncC-I1a chimeric plasmids and their significance as reservoirs of both antibiotic and metal resistance genes, reinforcing their epidemiological importance in the persistence and spread of multidrug-resistant *S*. Heidelberg.

### Chimeric IncC-I1a plasmids exhibit no detectable conjugal transfer activity

The annotation of the six *Salmonella* IncC-I1a plasmids revealed a partial deletion, present in all plasmids, affecting the approximately 28 Kb conserved region commonly found in IncC plasmids that encodes the majority of conjugal transfer genes^30^. This deletion resulted in the truncation of *traW* and the complete absence of *traU* and *traN*. Although the role of these IncC-related transfer genes has been inferred based on homology to corresponding genes in IncF plasmids and thus not experimentally confirmed in IncC, their importance for plasmid transfer is supported by these prior studies in IncF plasmids^31^. Specifically, deletion of *traW* in IncF plasmids has been shown to abolish conjugative transfer, while deletion of *traU* or *traN* significantly reduces transfer efficiency^31^.

To investigate the functional impact of this deletion on the six IncC-I1a plasmids, we assessed their ability to undergo conjugative transfer using pSH-363.78 as a representative. Solid mating experiments yielded no transconjugants, confirming either the lack of conjugative ability or transfer frequencies below the detection limit (≤1×10^−9^). Although IncC plasmids previously identified in *Salmonella* strains are often capable of conjugative transfer into new hosts^32^, our results align with earlier reports of such plasmids from other *S*. Heidelberg^33^ and *S*. Newport^32, 34^ isolates that lack this ability. These findings indicate that the observed deletions in key conjugal transfer genes likely account for the inability of these IncC-I1a plasmids to self-transfer, suggesting the loss of their conjugal function. However, the isolation of these nearly genetically similar IncC-I1a plasmids from *S*. Heidelberg strains belonging to three distinct PFGE-types^3^, together with the presence of predicted origins of transfer within the IncI1a integrated fragment (93,608 to 93,689 nt) and the IncC backbone (208,367 to 208,764 nt), suggests that these plasmids are mobilizable and therefore capable of horizontally spreading antibiotic resistance and virulence determinants.

### pSH-363.78 plasmid genetic structure confers high stability and marginal fitness cost

The stability of pSH-363.78 plasmid within a bacterial population was assessed by propagating *E. coli* DH10b::pSH-363.78 for approximately 90 generations with absence of selective pressure, indicating that pSH-363.78 was stably maintained in 100% of the bacterial cells per generation. This high stability suggests the presence of robust maintenance mechanisms that ensure persistence over time.

Similarly to other IncC plasmids, the IncC backbone of these chimeric plasmids harboured a type II toxin-antitoxin (TA) module, *tad* – *ata* (13,626 to 14283 nt), which has been previously experimentally validated to functions as an effective addiction module that maintains plasmid stability in an antibiotic-free environment^35^. In addition to this TA module, the plasmids also encode the type I *pndBCA* (85,490 to 85,754 nt) and type II *ccdAB* (114,121 to 114,646 nt) IncI1-associated TA modules^36^ as a result of acquiring an approximately 45-kb segment that is highly similar to an IncI1α-pST12 plasmid, as discussed above.

Due to this acquisition, these chimeric IncC-I1a plasmid harbour a panel of genes associated with plasmid maintenance, including : (i) the *cib* gene encoding the colicin Ib and its immunity gene *imm* (120,129 to 122,374 nt) that provide an advantage for the *Salmonella* host against competitors by inhibiting their growth particularly during inflammation^37^; (ii) the *ibfA* gene (116,928 to 117,272 nt) encoding a defence factor that confers abortive phage infection^38^; and (iii) the *klcA* gene (103,619 to 104,044 nt) encoding an anti-restriction protein that specifically inhibits endonuclease activity of type I restriction-modification systems^39^. In parallel, additionally to the native IncC parAB partitioning system (47,027 to 48,997 nt) and stbA partitioning protein (207,283 to 208,266 nt)^30^, these chimeric plasmids also harbour several genes predicted to encode partitioning-related proteins distributed throughout their sequence (13,305 to 13,598 nt; 25,307 to 26,245 nt; 99,357 to 101,315 nt; 110,297 to 111,681 nt; and 112,490 to 113,134 nt). Such a multi-layered strategy, involving multiple TA modules, partitioning systems, and other stability-associated genes, reflects an extensive evolutionary adaptation of these chimeric plasmids ensuring their stable persistence across diverse bacterial hosts.

Consistent with this extensive stabilization, the exponential growth rate of DH10b::pSH-363.78 (0,994; 95% CI 0.991-0.997) relative to DH10b was determined as an indicator of the fitness cost imposed by our chimeric plasmids on the host cell. Interestingly, carriage of pSH-363.78 imposed no significant fitness cost (p=0.056) under the conditions tested, contrary to the plasmid burden on bacterial hosts frequently reported in the literature^40^. In line with this, genomic analysis indicated the presence of the *psiAB* operon (98,152 to 99, 302 nt) and the *ssb* (101,665 to 102, 192 nt) gene, which encode anti-SOS factors^41, 42^, that prevent SOS-mediated stress responses that would otherwise impose burden in their bacterial host^43^. The absence of a measurable fitness cost together with the high stability of these plasmids imply that, under stewardship strategies that reduce selective pressures, such plasmids are unlikely to be purged by purifying selection and can persist as stable genetic scaffolds accumulating and retaining additional accessory traits. This, in turn, may facilitate the expansion of *S*. Heidelberg lineages carrying these chimeric plasmids, analogous to the pESI-like–associated global expansion of *S*. Infantis^44^.

### pSH-363.78 plasmid enhances bacterial pathogenicity

Sequence analysis of all six chimeric IncC-I1a plasmids indicated the presence of approximately 29 kb region encoding the *Yersinia* high-pathogenicity island (HPI; 155,142 to 184,942 nt), as previously described for the chimeric pESI plasmid in *S*. Infantis^9^. The involvement of this island in bacterial pathogenesis has been well demonstrated^45^, raising concerns about the contribution of these IncC-I1a plasmids in the virulence of the host bacterial hosts. To address this concern, the impact of harbouring the pSH-363.78 plasmid (as a representative of the six chimeric plasmids) on bacterial pathogenicity was evaluated using the *G. mellonella* invertebrate infection model. The LD_50_ value was determined to be 10^5^ CFU per larvae, and survival curves were compared between the isogenic *E. coli* strains (DH10b and DH10b::pSH-363.78). Our findings indicated that *E. coli* DH10b::pSH-363.78 exhibited significantly (*P =* < 0.05) higher killing rate over 96 h than *E. coli* DH10b or the PBS control, with mortality of 57% versus 27% and 0%, respectively (Figure 3). A significantly increased morbidity (via melanization score) was also observed in larvae infected with *E. coli* DH10b::pSH-363.78 over DH10b or the PBS control (Figure 4). Histopathological analysis of *G. mellonella* larvae six hours post infection further highlighted differences among the groups, with the highest levels of melanin deposition and haemocyte aggregation observed in larvae infected with DH10b::pSH-363.78 (Figure 5, Table S2). Visible alterations were also seen in bacteria adipose bodies among the three larvae groups, while gut morphology remained largely unaltered. Together these results indicate that the pSH-363.78 plasmid greatly enhances the overall virulence of its *E. coli* host in the *G. mellonella* invertebrate model, highlighting a role of these IncC-I1a plasmids in the pathogenicity of Enterobacteriaceae possibly owing to the presence of the *Yersinia* HPI. The convergence of multidrug resistance and virulence on a single mobile genetic element is increasingly documented and is recognized as a critical factor in the evolution of high-risk bacterial clones^46, 47^. Overall, the introduction of these plasmids in *S*. Heidelberg may drive the expansion of hypervirulent, multidrug-resistant lineages with increased zoonotic and epidemic potential.

**Figure 3.**
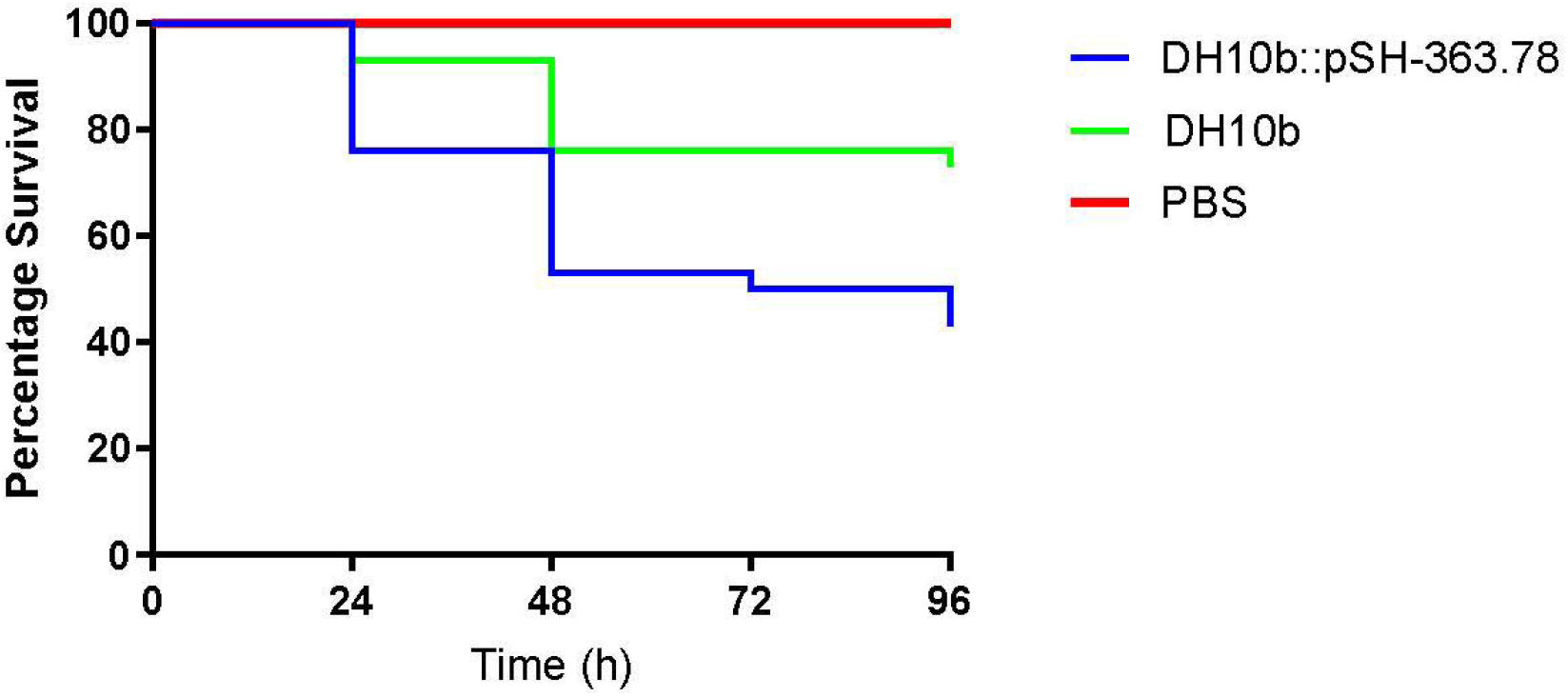
Live/dead assay of *Galleria mellonella* larvae infected with 10^5^ colony forming units/larvae of *E. coli* strains DH10b::pSH-363.78, DH10b or a phosphate buffered saline (PBS) control.

**Figure 4.**
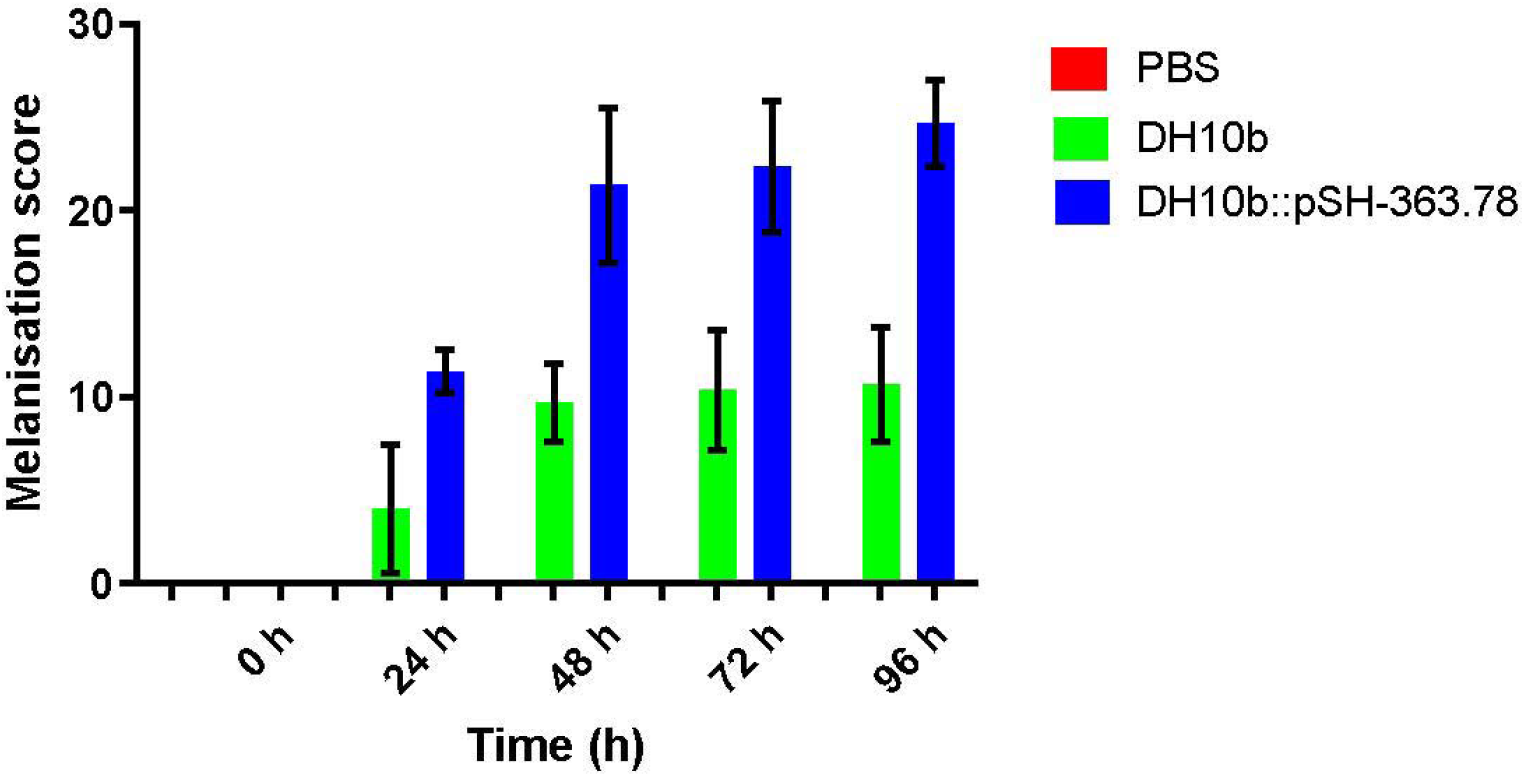
Melanization assay in *Galleria mellonella* larvae infected with 10^5^ colony forming units/larvae of *E. coli* strains DH10b::pSH-363.78, DH10b or a phosphate buffered saline (PBS) control.

**Figure 5.**
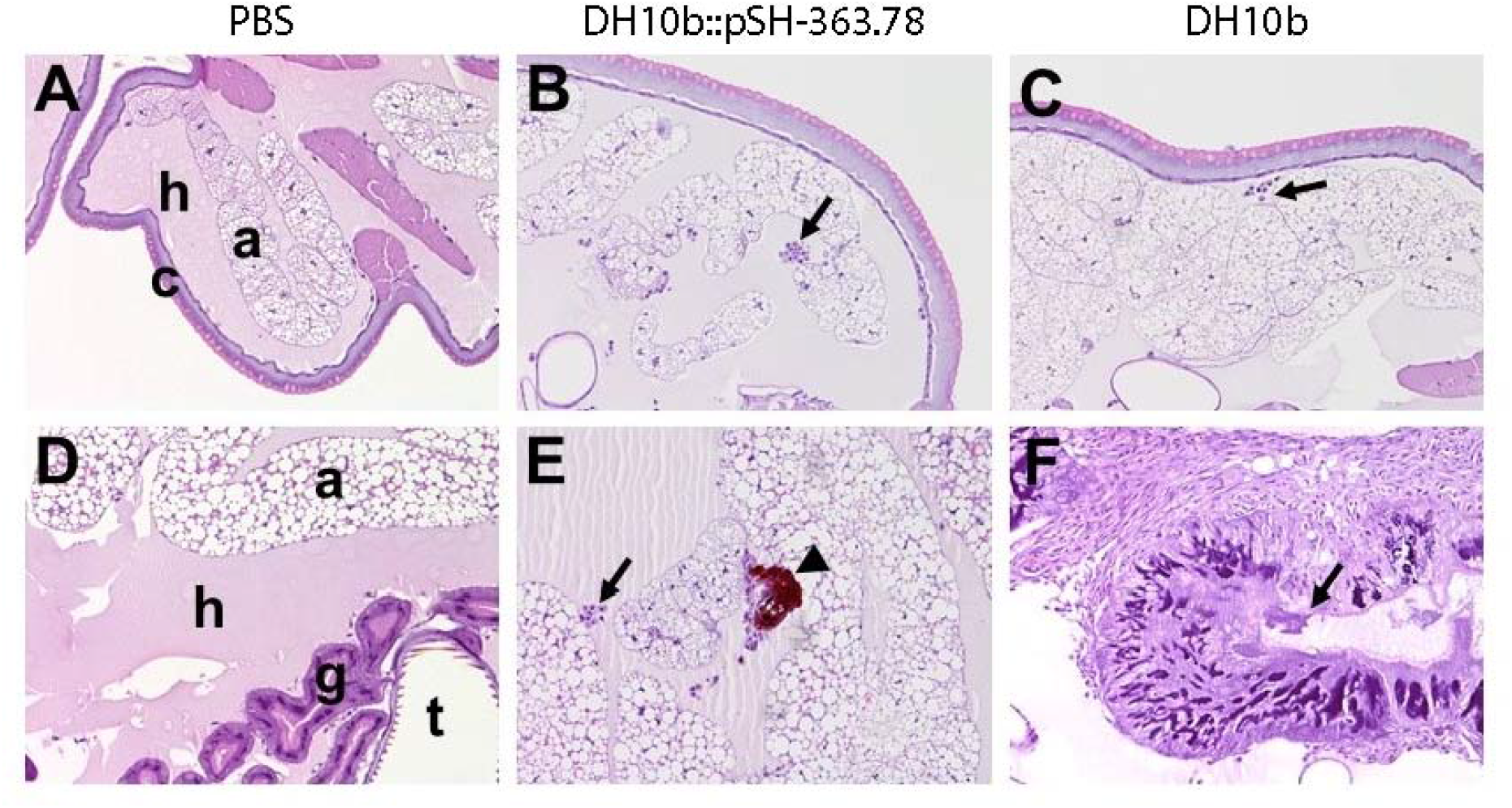
Histopathology of *Galleria mellonella* 6 h post inoculation with phosphate buffered saline (PBS), *E. coli* DH10b::pSH-363.78 (10^6^ colony forming units (CFU)/larvae and DH10b (10^6^ CFU/larvae). A. Normal structures from the larva anatomy: c= cuticle, h= haemolymph; a= adipose body. B-C. Clusters of haemocytes forming underneath the cuticle and surrounding adipose bodies (arrow). D. Normal structures from the larva anatomy: h= haemolymph; a= adipose body; g= gastrointestinal tract; t= trachea. E. Clusters of haemocytes observed surrounding adipose bodies (arrow) and melanization (arrowhead). F. Bacterial colonies observed within the gastrointestinal tract (arrow)

## CONCLUSIONS

The genetically novel chimeric IncC-I1a plasmids, as reported here, circulating among distinct PFGE-types of *S*. Heidelberg in the Netherlands display key functional and structural features reminiscent of pESI-like plasmids^9^. Specifically, these chimeric IncC-I1a plasmids combine multidrug resistance and virulence-associated determinants with accessory modules that confer high stability, efficient maintenance, and minimal fitness costs. Together, these features promote long-term persistence within bacterial populations even in the absence of antimicrobial selection pressure. This finding is of particular concern because similar plasmid–host associations have driven the global spread of pESI-positive *S*. Infantis in poultry production systems and throughout the food chain, contributing to the success of highly persistent and virulent multidrug-resistant lineages^44^. The emergence of functionally comparable IncC-I1a chimeras in *S*. Heidelberg suggests a similar evolutionary pathway, with an increased risk of sustained establishment in animal reservoirs and transmission to humans.

Overall, our findings indicate that these chimeric IncC-I1a plasmids act as powerful vehicles of adaptive evolution, promoting the convergence of antimicrobial resistance, virulence, and ecological fitness within a single mobile genetic element. This convergence might enhance bacterial adaptability and pathogenic potential while complicating control strategies that rely on the reduction of selective pressure. The introduction of such plasmids represents a growing threat to both public and animal health and reinforces the need for integrated genomic surveillance to monitor the dissemination of plasmids that couple antimicrobial resistance with enhanced virulence, as their emergence may complicate both treatment and containment of *Salmonella* outbreaks.

## Acknowledgements

We thank Prof. Dik J. Mevius for his assistance with initiating this project and Prof. Benno ter Kuile for providing the recipient strain *E. coli* MG1655::*yfp*.

## Transparency declarations

DC is currently employed by the European Food Safety Authority (EFSA). The positions and opinions presented in this article are those of the authors alone and are not intended to represent the views or scientific works of EFSA.

---

**Figure S1.**
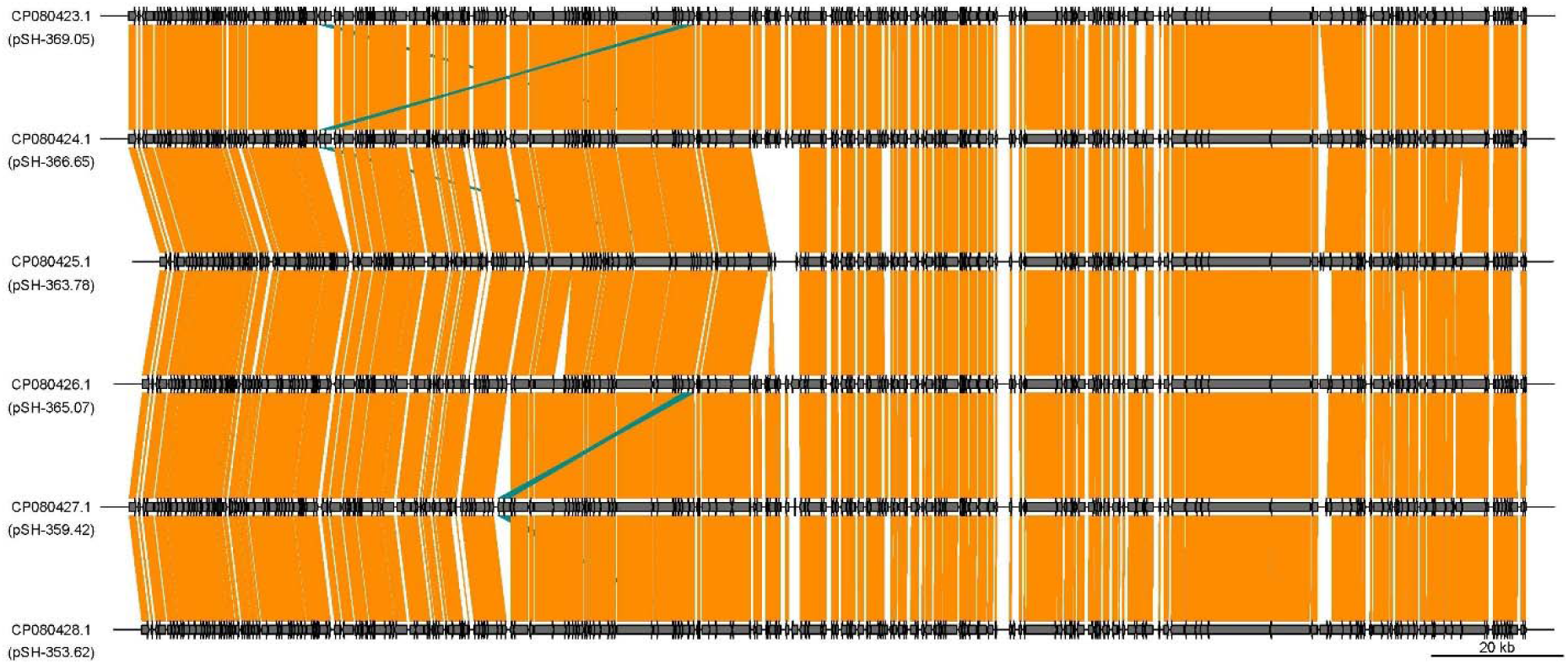
Genomic synteny comparison among the six novel chimeric IncC-I1a plasmids, generated using a custom R script with the *genoPlotR* package, represented by horizontal grey bars. Homologous regions between adjacent plasmids are highlighted by coloured blocks: orange blocks connect regions oriented on the same strand, while teal (blue-green) blocks connect regions in an inverted orientation. The intensity of the orange and teal colours indicates the level of sequence identity, with darker shades representing higher identity (up to 100%). Only homologous regions with at least 90% sequence coverage are shown.

**Table S1.** IncC-I1a plasmids included in this study and their characteristics.

| Plasmid ID | Host | Host source | Resistance gene(s) | Size (bp) | Accession number |
| --- | --- | --- | --- | --- | --- |
| pSH-353.62 | *S. Heidelberg* (Xbal.X) | PM | *bla*<sub>CMY-2</sub>, *tetA*, *sul2* | 212,338 | CP080428.1 |
| pSH-359.42 | *S. Heidelberg* (Xbal.1973) | PM | *bla*<sub>CMY-2</sub>, *tetA*, *sul2* | 214,320 | CP080427.1 |
| pSH-363.78 | *S. Heidelberg* (Xbal.1973) | PM | *bla*<sub>CMY-2</sub>, *tetA*, *sul2* | 209,580 | CP080425.1 |
| pSH-365.07 | *S. Heidelberg* (Xbal.1973) | PM | *bla*<sub>CMY-2</sub>, *tetA*, *sul2* | 212,341 | CP080426.1 |
| pSH-366.65 | *S. Heidelberg* (Xbal.1965) | FPA | *bla*<sub>CMY-2</sub>, *tetA*, *sul2* | 214,418 | CP080424.1 |
| pSH-369.05 | *S. Heidelberg* (Xbal.1973) | FPA | *bla*<sub>CMY-2</sub>, *tetA*, *sul2* | 214,427 | CP080423.1 |

FPA: food-producing animals, PM: poultry meat.

**Table S2.**
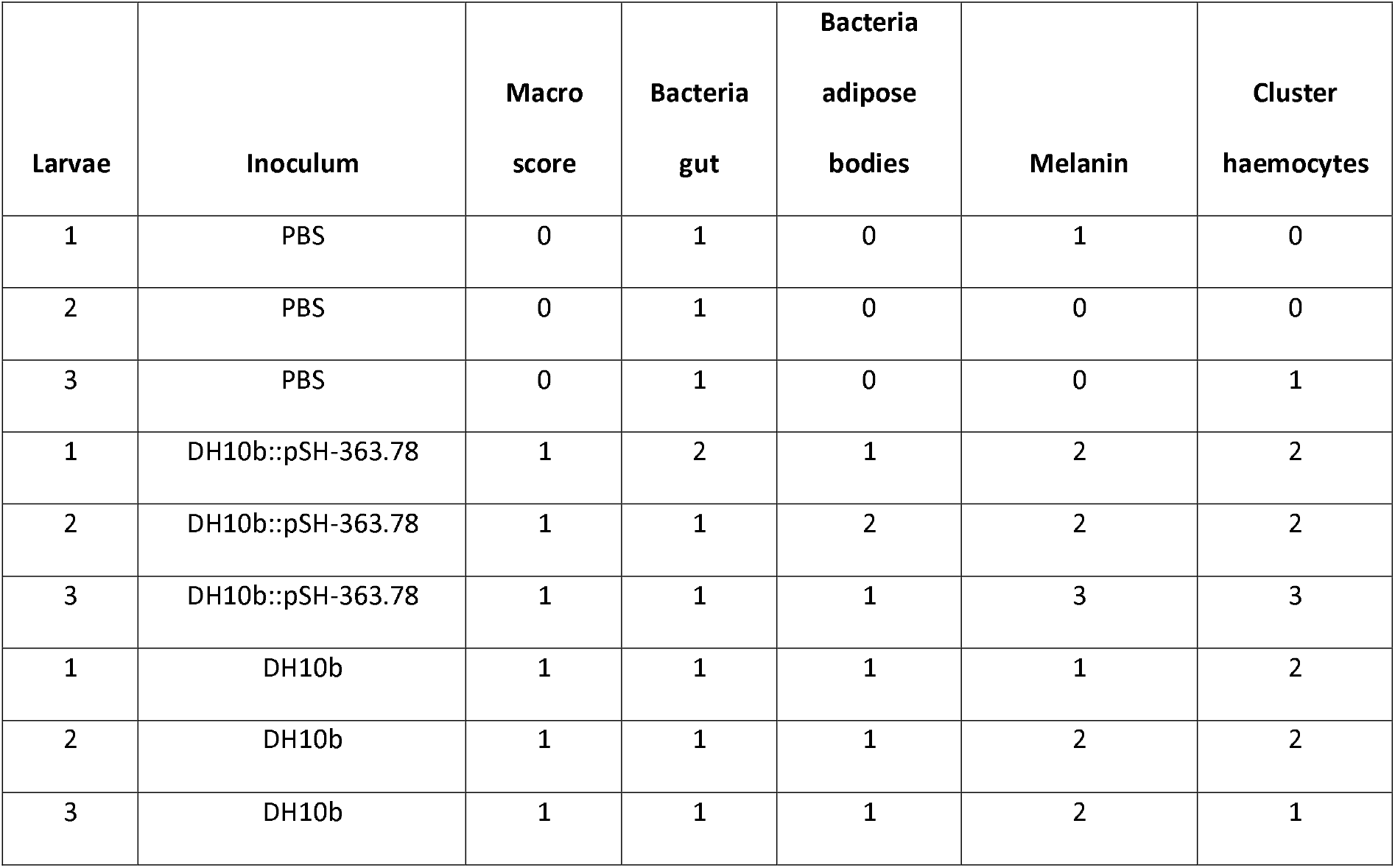
Histopathology scores for *Galleria mellonella* 6 h post inoculation with PBS, *E. coli* DH10b::pSH-363.78 (10^6^ colony forming units (CFU)/larvae and DH10b (10^6^ CFU/larvae).

| Larvae | Inoculum | Macro<br>score | Bacteria<br>gut | Bacteria<br>adipose<br>bodies | Melanin | Cluster<br>haemocytes |
| --- | --- | --- | --- | --- | --- | --- |
| 1 | PBS | 0 | 1 | 0 | 1 | 0 |
| 2 | PBS | 0 | 1 | 0 | 0 | 0 |
| 3 | PBS | 0 | 1 | 0 | 0 | 1 |
| 1 | DH10b::pSH-363.78 | 1 | 2 | 1 | 2 | 2 |
| 2 | DH10b::pSH-363.78 | 1 | 1 | 2 | 2 | 2 |
| 3 | DH10b::pSH-363.78 | 1 | 1 | 1 | 3 | 3 |
| 1 | DH10b | 1 | 1 | 1 | 1 | 2 |
| 2 | DH10b | 1 | 1 | 1 | 2 | 2 |
| 3 | DH10b | 1 | 1 | 1 | 2 | 1 |

## References

1. Public Health Agency of Canada. National Enteric Surveillance Program (NESP), Annual Summary 2012.) (2014).

2. CDC. National Antimicrobial Resistance Monitoring System for Enteric Bacteria (NARMS): Human Isolates Final Report, 2012.). Department of Health and Human Services (2014).

3. Liakopoulos A, et al. Extended-Spectrum Cephalosporin-Resistant Salmonella enterica serovar Heidelberg Strains, the Netherlands(1). Emerg Infect Dis 22, 1257–1261 (2016).

4. Hoffmann M, et al. Comparative genomic analysis and virulence differences in closely related salmonella enterica serotype heidelberg isolates from humans, retail meats, and animals. Genome Biol Evol 6, 1046–1068 (2014).

5. Burt CR, Proudfoot JC, Roberts M, Horowitz RH. Fatal myocarditis secondary to Salmonella septicemia in a young adult. J Emerg Med 8, 295–297 (1990).

6. Figueroa-Bossi N, Bossi L. Inducible prophages contribute to Salmonella virulence in mice. Molecular microbiology 33, 167–176 (1999).

7. Rice PA, Craven C, Wells JG. Salmonella heidelberg enteritis and bacteremia. An epidemic on two pediatric wards. Am J Med 60, 509–516 (1976).

8. Cohen E, Rahav G, Gal-Mor O. Genome Sequence of an Emerging Salmonella enterica Serovar Infantis and Genomic Comparison with Other S. Infantis Strains. Genome Biol Evol 12, 151–159 (2020).

9. Aviv G, et al. A unique megaplasmid contributes to stress tolerance and pathogenicity of an emergent Salmonella enterica serovar Infantis strain. Environ Microbiol 16, 977–994 (2014).

10. Wick RR, Judd LM, Gorrie CL, Holt KE. Unicycler: Resolving bacterial genome assemblies from short and long sequencing reads. PLoS Comput Biol 13, e1005595 (2017).

11. Overbeek R, et al. The SEED and the Rapid Annotation of microbial genomes using Subsystems Technology (RAST). Nucleic Acids Res 42, D206–214 (2014).

12. Zankari E, et al. Identification of acquired antimicrobial resistance genes. J Antimicrob Chemother 67, 2640–2644 (2012).

13. Carattoli A, et al. In silico detection and typing of plasmids using PlasmidFinder and plasmid multilocus sequence typing. Antimicrob Agents Chemother 58, 3895–3903 (2014).

14. Siguier P, Perochon J, Lestrade L, Mahillon J, Chandler M. ISfinder: the reference centre for bacterial insertion sequences. Nucleic Acids Res 34, D32–36 (2006).

15. Ambrose SJ, Harmer CJ, Hall RM. Evolution and typing of IncC plasmids contributing to antibiotic resistance in Gram-negative bacteria. Plasmid 99, 40–55 (2018).

16. Liakopoulos A, et al. Genomic and functional characterisation of IncX3 plasmids encoding bla(SHV-12) in Escherichia coli from human and animal origin. Sci Rep 8, 7674 (2018).

17. Letunic I, Bork P. Interactive Tree of Life (iTOL) v6: recent updates to the phylogenetic tree display and annotation tool. Nucleic Acids Res 52, W78–W82 (2024).

18. Li Y, et al. PlasmidScope: a comprehensive plasmid database with rich annotations and online analytical tools. Nucleic Acids Res 53, D179–D188 (2025).

19. Gu Z, Gu L, Eils R, Schlesner M, Brors B. circlize Implements and enhances circular visualization in R. Bioinformatics 30, 2811–2812 (2014).

20. Guy L, Kultima JR, Andersson SG. genoPlotR: comparative gene and genome visualization in R. Bioinformatics 26, 2334–2335 (2010).

21. Carattoli A, Bertini A, Villa L, Falbo V, Hopkins KL, Threlfall EJ. Identification of plasmids by PCR-based replicon typing. J Microbiol Methods 63, 219–228 (2005).

22. Liakopoulos A, et al. Genomic and functional characterisation of IncX3 plasmids encoding blaSHV-12 in Escherichia coli from human and animal origin. Sci Rep 8, 7674 (2018).

23. Betts JW, Hornsey M, Wareham DW, La Ragione RM. In vitro and In vivo Activity of Theaflavin-Epicatechin Combinations versus Multidrug-Resistant Acinetobacter baumannii. Infect Dis Ther 6, 435–442 (2017).

24. Wang-Kan X, et al. Lack of AcrB Efflux Function Confers Loss of Virulence on Salmonella enterica Serovar Typhimurium. mBio 8, (2017).

25. Senior NJ, et al. Galleria mellonella as an infection model for Campylobacter jejuni virulence. J Med Microbiol 60, 661–669 (2011).

26. Hoffmann M, et al. Complete Genome Sequence of a Multidrug-Resistant Salmonella enterica Serovar Typhimurium var. 5-Strain Isolated from Chicken Breast. Genome Announc 1, (2013).

27. Labbe G, et al. Complete Genome and Plasmid Sequences of Three Canadian Isolates of Salmonella enterica subsp. enterica Serovar Heidelberg from Human and Food Sources. Genome Announc 4, (2016).

28. Allain M, et al. IncC plasmid genome rearrangements influence the vertical and horizontal transmission tradeoff in Escherichia coli. Antimicrob Agents Chemother 68, e0055424 (2024).

29. Pal C, Bengtsson-Palme J, Kristiansson E, Larsson DG. Co-occurrence of resistance genes to antibiotics, biocides and metals reveals novel insights into their co-selection potential. BMC Genomics 16, 964 (2015).

30. Harmer CJ, Hall RM. The A to Z of A/C plasmids. Plasmid 80, 63–82 (2015).

31. Kishida K, et al. Contributions of F-specific subunits to the F plasmid-encoded type IV secretion system and F pilus. Mol Microbiol 117, 1275–1290 (2022).

32. Welch TJ, et al. Multiple antimicrobial resistance in plague: an emerging public health risk. PLoS One 2, e309 (2007).

33. Han J, et al. DNA sequence analysis of plasmids from multidrug resistant Salmonella enterica serotype Heidelberg isolates. PLoS One 7, e51160 (2012).

34. Poole TL, Edrington TS, Brichta-Harhay DM, Carattoli A, Anderson RC, Nisbet DJ. Conjugative transferability of the A/C plasmids from Salmonella enterica isolates that possess or lack bla(CMY) in the A/C plasmid backbone. Foodborne Pathog Dis 6, 1185– 1194 (2009).

35. Qi Q, Kamruzzaman M, Iredell JR. The higBA-Type Toxin-Antitoxin System in IncC Plasmids Is a Mobilizable Ciprofloxacin-Inducible System. mSphere 6, e0042421 (2021).

36. Smith H, et al. Characterization of epidemic IncI1-Igamma plasmids harboring ambler class A and C genes in Escherichia coli and Salmonella enterica from animals and humans. Antimicrob Agents Chemother 59, 5357–5365 (2015).

37. Nedialkova LP, et al. Inflammation fuels colicin Ib-dependent competition of Salmonella serovar Typhimurium and E. coli in enterobacterial blooms. PLoS Pathog 10, e1003844 (2014).

38. Hiley L, Graham RMA, Jennison AV. Characterisation of IncI1 plasmids associated with change of phage type in isolates of Salmonella enterica serovar Typhimurium. BMC Microbiol 21, 92 (2021).

39. Goryanin, II, Kudryavtseva AA, Balabanov VP, Biryukova VS, Manukhov IV, Zavilgelsky GB. Antirestriction activities of KlcA (RP4) and ArdB (R64) proteins. FEMS Microbiol Lett 365, (2018).

40. Baltrus DA. Exploring the costs of horizontal gene transfer. Trends in ecology & evolution 28, 489–495 (2013).

41. Petrova V, Chitteni-Pattu S, Drees JC, Inman RB, Cox MM. An SOS inhibitor that binds to free RecA protein: the PsiB protein. Mol Cell 36, 121–130 (2009).

42. Jones AL, Barth PT, Wilkins BM. Zygotic induction of plasmid ssb and psiB genes following conjugative transfer of Incl1 plasmid Collb-P9. Mol Microbiol 6, 605–613 (1992).

43. Curtsinger HD, Martinez-Absalon S, Liu Y, Lopatkin AJ. The metabolic burden associated with plasmid acquisition: An assessment of the unrecognized benefits to host cells. Bioessays 47, e2400164 (2025).

44. Alba P, et al. Molecular epidemiology of Salmonella Infantis in Europe: insights into the success of the bacterial host and its parasitic pESI-like megaplasmid. Microb Genom 6, (2020).

45. Carniel E. The Yersinia high-pathogenicity island: an iron-uptake island. Microbes Infect 3, 561–569 (2001).

46. Yang X, Wai-Chi Chan E, Zhang R, Chen S. A conjugative plasmid that augments virulence in Klebsiella pneumoniae. Nat Microbiol 4, 2039–2043 (2019).

47. Linkevicius M, et al. Cross-border spread of a mosaic resistance (OXA-48) and virulence (aerobactin) plasmid in Klebsiella pneumoniae: a European Antimicrobial Resistance Genes Surveillance Network investigation, Europe, February 2019 to October 2024. Euro Surveill 30, (2025).

